# Context-dependence of deterministic and nondeterministic contributions to closed-loop steering control

**DOI:** 10.1101/2024.07.26.605325

**Authors:** Seth W. Egger, Sander W. Keemink, Mark S. Goldman, Kenneth H. Britten

## Abstract

In natural circumstances, sensory systems operate in a closed loop with motor output, whereby actions shape subsequent sensory experiences. A prime example of this is the sensorimotor processing required to align one’s direction of travel, or heading, with one’s goal, a behavior we refer to as steering. In steering, motor outputs work to eliminate errors between the direction of heading and the goal, modifying subsequent errors in the process. The closed-loop nature of the behavior makes it challenging to determine how deterministic and nondeterministic processes contribute to behavior. We overcome this by applying a nonparametric, linear kernel-based analysis to behavioral data of monkeys steering through a virtual environment in two experimental contexts. In a given context, the results were consistent with previous work that described the transformation as a second-order linear system. Classically, the parameters of such second-order models are associated with physical properties of the limb such as viscosity and stiffness that are commonly assumed to be approximately constant. By contrast, we found that the fit kernels differed strongly across tasks in these and other parameters, suggesting context-dependent changes in neural and biomechanical processes. We additionally fit residuals to a simple noise model and found that the form of the noise was highly conserved across both contexts and animals. Strikingly, the fitted noise also closely matched that found previously in a human steering task. Altogether, this work presents a kernel-based analysis that characterizes the context-dependence of deterministic and non-deterministic components of a closed-loop sensorimotor task.

**New and noteworthy:** We use nonparametric systems identification techniques to assess the context-dependence of deterministic and nondeterministic contributions to a closed-loop behavior. Classical approaches assume a fixed transformation between sensory input and motor output. Here, we reveal strong changes to the measured sensorimotor transformations with behavioral context. In contrast, noise within the transformation exhibited a consistent form across contexts, subjects, and species. Together, this work demonstrates how context affects the systematic and stochastic components of a closed-loop behavior.

## 1. Introduction

The neural machinery for sensation and motor control are often thought of as distinct, separable systems. This has led to experimental approaches that isolate each – experiments to study sensory systems typically utilize preparations that only require the animal to sit quietly as their sensory epithelia are stimulated (1–4) and those to study the motor system use reduced sensory inputs to focus on the control of the effector (5–7). While this approach has been extremely fruitful, neural systems evolved to operate in a regime in which sensory and motor systems drive each other in a closed-loop – sensory input drives motor output, which modifies sensory input and subsequent motor output. As a result, many of the conclusions reached about sensory processing or motor control may not hold during the natural operation of neural systems. Indeed, neurophysiological experiments combining sensory stimulation with active movements suggest that neural responses in areas typically associated with sensory processing are impacted by ongoing motor behavior (2, 8–16).

While neural systems may be optimized to operate in a closed-loop regime, classic studies of sensorimotor responses have been done in the open-loop regime where the experimenter controls the stimulus, enabling tight regulation of behavior. By contrast, in closed-loop systems, motor errors drive responses. Such motor errors reflect both sensorimotor neural processing and also mechanical features of the musculoskeletal system, such as viscous drag and spring-like forces, that shape the speed and amplitude of movements (17–19). For example, to steer toward a distant target, humans control their direction of travel in real time by comparing the direction of locomotion to the visual direction of the target (20). This provides an error signal that cues the direction and magnitude of the movements made to reach the target (21–23). Therefore, the components of the system driving a response are tightly correlated, making the open-loop sensorimotor transformations occurring within the nervous system and motor effectors difficult to discern from the observed closed-loop response.

Classically, these challenges were addressed by modeling the closed-loop response with linear systems chosen from a family of functions with straightforward interpretations (24). In the context of steering control, this approach leads to models with a proportional response to the steering error, and potentially its derivative or integral, as well as terms that model the physical constraints on the appendages and actuators controlling steering output (25–29). Implicit in most models is the assumption that the system is linear and that temporal dynamics of the response arise heavily from the mechanical properties of the motor system. There are three ways in which these assumptions might lead to erroneous identification of the properties of the underlying steering system. First, the components of the response that are assumed to be mechanical in origin can often be explained by the filtering performed by sensory or sensorimotor processing (24, 30). Second, because the motor effectors are often modeled as stationary over time and conditions, changes in steering behavior are assumed to stem from changes in sensorimotor processing (23, 26, 29). However, these mechanical properties flexibly adjust according to behavioral context (31) and therefore may also contribute to the observed changes. Finally, residual steering output not explained by the model is typically attributed to noise in the neural system, but substantial components of the residual behavior might simply be missed by the model. For example, while most models assume a linear system, sensorimotor systems have substantial nonlinear properties that might contribute to behavior in significant ways (32). We therefore set out to determine the degree to which nonlinearities, noise, and behavioral context contribute to steering behavior with a novel method with more limited assumptions.

More recent approaches to systems identification instead have specified a broad space of potential transformation functions through a nonparametric basis and then used observed responses to select for the transformation that best predicts the data (33–37). We refer to these approaches as kernel-based methodologies. A strength of these approaches is that they afford the experimenter confidence about the inferences made from the model fit to the data. For example, it is possible to formulate the basis such that it spans the space of all possible linear models. In this case, the experimenter can be confident that the linear components of the transformation function are captured by the model and any behavior unaccounted for must come from either nonlinearities or noise in the transformation function. When combined with trial-averaging to remove noise, these approaches provide a powerful means to place limits on the degree to which the transformation can be considered nonlinear (38, 39).

Recent work has extended kernel-based methods to closed-loop sensorimotor systems (40–42). We therefore set out to apply this approach to measure the contributions of linearities, nonlinearities, and noise to steering behavior. We adapted nonparametric kernel methods for systems identification to a steering task that required the monkey to manipulate a joystick to control its instantaneous angular velocity in a virtual environment to match its direction of travel with a distant target (22). Application of our kernel method found that a linear model describes a large fraction of the steering behavior of macaques. Further, the form of kernel identified with our nonparametric approach was consistent with those proposed by previous parametric model-based methods, validating this approach to identifying sensorimotor transformations (25–27, 29).

However, our method also revealed several new features of steering behavior. First, we found that, contrary to the assumptions of previous models, the components of the steering system commonly modeled as constant physical constraints associated with the motor system changed with experimental context. Second, much of the trial-by-trial variations in steering response remained unexplained after accounting for the linear portion of the transformation. The statistics of this residual behavior were strikingly similar across paradigms and monkeys. Application of a simple noise model captured the statistics of the residual behavior remarkably well, suggesting the interpretation that unexplained behavioral variance arises from noise in sensory processing in a manner analogous to other sensorimotor behaviors (43, 44). Overall, we demonstrate that our kernel-based approach allows us to tease apart the influence of linear versus nonlinear and noise contributions to steering behavior and provides a framework for modeling sensorimotor transformations in closed-loop designs.

## 2. Methods

### 2.1. Animals

We trained two adult female rhesus macaques to manipulate a joystick to steer towards a target by operant conditioning techniques. For details on training, see (22). All experiments were conducted with the approval of the UC Davis Animal Care and Use Committee and adhered to ILAR and USDA guidelines for the treatment of experimental animals.

### 2.2. Apparatus

Stimuli were generated on a dedicated computer by custom software (written by A. L. Jones and D. J. Sperka) that used OpenGL libraries running under a real-time Linux kernel. The display computer ran at a resolution of 1024x768 pixels at 85 Hz. Both monkeys viewed the CRT monitor (Mitsubishi Diamond Pro 21) at a distance of 28 cm, so that the monitor subtended 60 deg horizontally by 45 deg vertically. The maximum luminance of the monitor was set to 60 cd/m^2^. The display computer received commands from an experimental computer running Rex, the NIH public domain package. An analog voltage signal from the joystick, sampled at 1 kHz with a 12-bit analog-to-digital converter by the experimental computer, controlled the angular velocity of the animal’s trajectory in the virtual world. This signal was sampled at 85 Hz for the purpose of updating the next frame of the display computer. We set the gain of the joystick to 255 deg/s and 85 deg/s at maximum stick deflection for monkeys F and J, respectively. We chose the gain based on behavioral performance early in training such that each animal’s joystick deflections were linear with respect to joystick output. For additional details on the hardware and software used, see (22).

### 2.3. Behavioral task

The monkeys sat in a darkened room with their views centered on the display monitor. We displayed a simulated environment that consisted of a distant red target and a dotted ground plane under a dark sky. Dots had a luminance of 60 cd/m^2^ and the ground plane was 7 cd/m^2^. We moved the dots on the ground plane so that the global pattern of motion simulated a translational movement aligned with the monkey’s field of view at a constant speed of 2.13 m/s at a height of 50 cm. See Figure 1a for an example of the scene displayed to the monkey. At the beginning of a trial, the target appeared a few degrees from the center of the screen and the ground plane began to move. Each monkey manipulated a single-axis joystick with its right hand, wrist and arm to control the direction of movement across the ground plane; movements of the joystick resulted in a turn with an angular velocity proportional to the stick displacement. Maximum deflections of the joystick were on the order of 5 cm. Figure 1b provides a schematic of a single frame of the experiment from an overhead view. In this example, the target (*T*) and the heading (*H*) directions do not agree. This results in the monkey observing a steering error (*x*) of (*T* −*H*) deg. In response to this error, the monkey makes an appropriate movement with the joystick (left in this example), changing the heading to better match the target direction and decrease the error. Figures 1c and 1d provide example traces of the target position (red), heading (blue), steering error (purple) and monkey responses (black) from two experimental contexts, referred to as the step and drift contexts, respectively.

**Figure 1.**
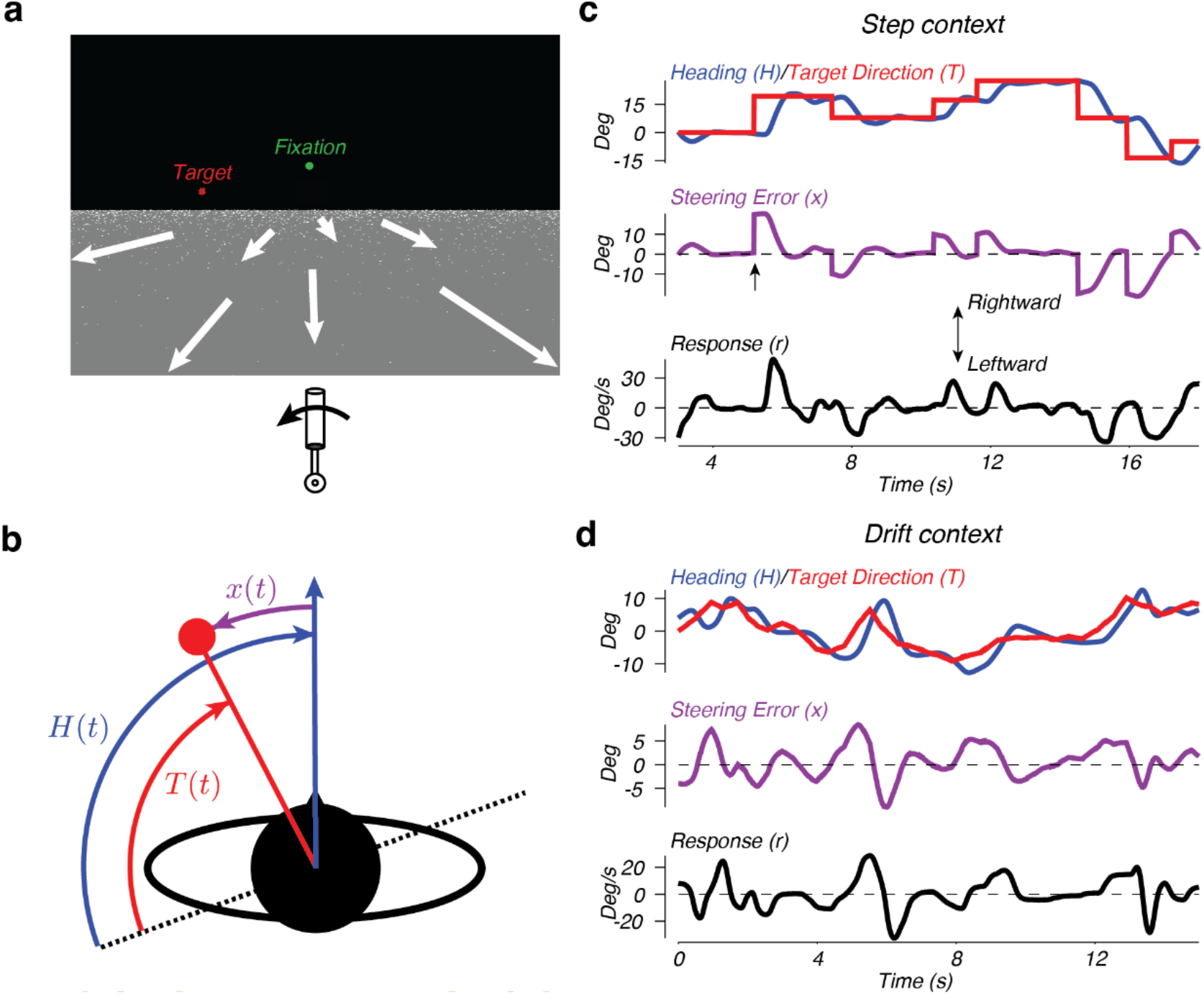
Design of the steering task. a) View from the cockpit. The target is represented by the red dots. The green spot represents the fixation point (drift context only). The joystick and arrow below the scene represent the correct steering behavior for this example. The white arrows illustrate the movement of dots on the ground plane and were not visible to the monkey. b) Overhead view of the experiments. The red dot represents the target and the blue arrow represents the current direction of travel of the monkey. The dotted line represents the arbitrary reference frame in which the target position (*T*) and the heading (*H*) were measured. *x* represents the steering error. c) An example of 15 s of steering in the step context. Top: target direction *T*(*t*) (red trace) and heading *H*(*t*) (blue trace), in world coordinates. Middle: steering error *x*(*t*). Bottom: steering response, *r*(*t*). d) An example of 15 s of steering in the drift context. Color conventions are the same as in panel c.

#### 2.3.1. Step context

In the step context (Figure 1c), the target remained stationary in world coordinates for periods of several seconds before randomly stepping to a new location. The time between steps was chosen from a truncated exponential distribution (1000 ms minimum, 2000 ms on average). The amplitude of each step was chosen so that the resulting steering error would be 5, 10, 15, 20, or 25 deg in amplitude. The probability of the occurrence of a step decreased with amplitude, but the range of step sizes varied from day to day. The target was a solid red disc 0.25 deg in diameter. Each trial began with the target located centrally and the ground plane stationary for 500 ms before the target stepped between 5 and 25 deg to the left or right, the ground plane began to move, and steering could begin. An example trace from this paradigm shows that the target location in world coordinates exhibits large steps followed by stationary periods (Figure 1c, red trace). Each trial lasted 15-30 s with a 2 s intertrial interval. For these experiments, the monkeys could move their eyes at will, and reliably tracked the steering target with their gaze. For a detailed analysis of the behavior in the step context, see (22).

#### 2.3.2. Drift context

In the second class of experiments, the target slowly drifted in the world at random speeds (Figure 1d). In these experiments, the target was 20 red pixels selected at random from locations within 0.25 deg of the true target location. The location of the red pixels within the disc changed randomly at a mean rate of 1 per frame. Additionally, unlike the first class of experiments, the monkeys were required to maintain their gaze within 3 degrees of a green fixation point located 10 deg above the center of the screen. Each trial began by displaying the fixation point alone on the screen. After the monkey fixated for 150 ms, the target and ground plane appeared but remained stationary for an additional 500 ms. At this point the target moved to a new location 4 deg to the left or right, the ground plane started to move, and steering could begin. For the rest of a 15 s trial, the monkeys were required to continue to fixate while steering to the target. The target moved in the world at an angular velocity chosen from a zero mean Gaussian with 0.1 deg/s standard deviation. Every 259-494 ms a new velocity was chosen from the same distribution, resulting in a random drift through the simulated world. As a result, the monkeys needed to constantly steer to receive reward. Two additional manipulations also occurred. First, zero mean noise with 0.1 deg standard deviation was added to the displayed location of the target every 94 ms. Additionally, the displayed heading across the ground plane was also corrupted by zero mean noise with 2.5 deg standard deviation, updated every 94 ms. The monkeys were rewarded immediately upon achieving a heading direction within 3 deg of the target direction and at a rate that increased with the duration the target was maintained within 3 deg. The reward function was based on the location of the target without noise relative to the heading without noise.

### 2.4. Steering model

A primary goal of this work was to identify the model that best captures the linear portion of the response, *r*(*t*), to the past series of observed steering errors. We modeled the monkey steering system as a linear function of a noisy estimate of the steering error, *x*(*t*) +*n*(*t*),

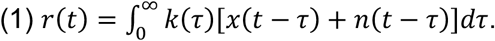

*k*(*τ*) describes the pattern of weights given to past steering errors for the linear response. In this paper, we refer to *k*(*τ*) as the linear kernel of the steering system. The deterministic portion of the response, 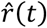, is the weighted sum of the recent steering errors:

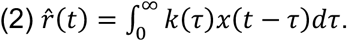

The stochastic portion of the response, *q*(*t*), is the weighted sum of the history of noise:

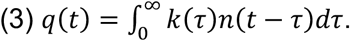

Because steering in a closed-loop induces significant autocorrelation in the steering error, typical regression-based methods identify kernels with non-causal components. To avoid these artifacts, we used a set of basis functions that only span the causal time lags to describe the kernel. The kernel is modeled as the weighted sum of these basis functions:

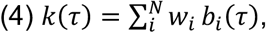

where *w*_*i*_ is the weight given to the corresponding basis function, *b*_*i*_(*τ*), and *N* is the total number of basis functions. We chose our basis functions as a set of overlapping cosine bumps defined as:

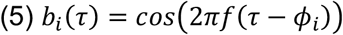

for −*π*/2 < 2*πf* (*τ* −*ϕ*_*i*_) < *π*/2 and 0 otherwise. For subsequent analysis in the text, we chose *N* = 56 basis functions with the centers of each half cosine bump, *ϕ*_*i*_, linearly sampling the possible lags between 0.125 and 4.708 s, and the frequency of each bump, *f* = 2 Hz. Our choice of basis functions constrains the possible kernels to a subspace of linear models that are relatively smooth, have a finite memory, and are forced to be zero at lags less than or equal to zero. To arrive at this set of constraints, we tested several different forms of bases including triangular functions and half-cosine bumps that linearly tile compressed time (e.g. *b*_*i*_(*τ*) =cos(2*π*(*g*[*ψτ*] −*ϕ*_*i*_)*>*, where *g*[*x*] is a square-root or logarithmic function). We further tested bases of different widths and spacing. In general, the exact form of the basis function, width, and spacing did not qualitatively change any results and the differences in predictive performance were on the order of 1% variance explained. The only exception to this were basis functions that approached delta functions, which lead to kernel functions with the majority of weighting given to impossibly short time lags. This, combined with a comparison of the results to those using a standard parametric model (second-order linear model; see below), suggests our basis set covered the linear subspace containing the steering system, up to a constraint on the abruptness of the onset response, which is forced to be somewhat smooth.

We fit the values of the basis function weights, *w*_*i*_, by minimizing the sum of the squared errors between the predicted responses, 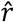, and the observed responses. To avoid overfitting, we split our data into training and validation sets. We uniformly sampled (without replacement) 100 trials for fitting and used the remaining trials for validation. In total, we had 728 and 446 total trials for monkeys F and J, respectively, from the drift context and 3908 and 2133 total trials for F and J, respectively, from the step context. The results did not depend substantially on the subset of trials used for training the model or the number of trials used for training. Finally, to remove artifacts due to the asymmetric overlap in the final 2 basis functions relative to the others, we set the amplitude of the kernel to zero for lags greater than 4.835 s. To test our model, we provided only the initial heading and the target position in world coordinates for the duration of the experiment.

### 2.5. Residuals analysis

The analysis of responses to single steps of the target reveal that the monkey steering system deviates significantly from the mean on a trial by trial basis (22). To assess the source of the residual behavior in monkey steering responses, we compared the residual spectrum observed from the data to the spectrum of residuals expected based on simulations of the kernel with noise. The residual steering behavior not explained by the linear model was calculated as 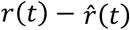. We calculated the power spectrum as the squared magnitudes of the Fourier coefficients calculated by a fast Fourier transform of the residuals for each trial and averaged across trials to find the average residual spectrum. We then normalized the resulting spectrum by the total power across frequencies.

*Multiplicative noise model*. Equation (1) describes a “multiplicative” noise model, where the noise term is multiplied by the sensorimotor kernel, *k*(*τ*). To assess the ability of the model to explain the average residual spectra, we ran simulations of the model and compared these simulations to the experimental data. We used the target position and initial heading in world coordinates from the actual experiments as the initial conditions for each simulated monkey and experiment. To simulate multiplicative noise, we added zero-mean, independent Gaussian noise to the error signal before passing it to the linear kernel. We then found the residual spectrum using the same procedure as for the actual steering responses. We chose the standard deviation of the simulated noise to be 1 deg/s, but the results after normalizing each spectrum by the total variance in steering error did not depend on the level of noise simulated.

*Estimation of the noise spectrum*. To empirically estimate the spectrum of the noise input, Φ_*nn*_, added to the steering error by the monkey, we measured the power spectra of the target position, Φ_*TT*_, the systematic response from the linear kernel, 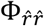, and the residuals, Φ_*qq*_, and used these quantities to estimate the spectrum of the noise as:

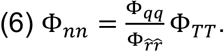

Assuming the noise is independent of the target position and the animal is accurately modeled by a linear system, this equation will find the spectrum of the noise added to the steering error during error estimation (45) (see Appendix for derivation). All power spectra were calculated in the same manner as described above for the residual spectra.

### 2.6. Second-order linear model

Previous models of steering in humans have typically used a second-order linear system (23, 25, 26, 29). In such models, accelerations of the hand controlling the joystick are determined by the steering error in the recent past, minus terms for the velocity and position of the hand:

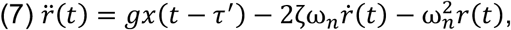

where *g* is the weight given to the steering error *τ*′ seconds in the past. The second term of equation (7) represents resistance to motion by viscous drag-like forces. The last term represents resistance to nonzero position through a spring-like restoring force. The parameters ζ and ω_n_ control the stiffness and viscosity of the system. The damping ratio, ζ, determines the level of damping in the system, while the undamped natural (angular) frequency ω_*n*_ controls the frequency of oscillation. Converting to the Laplace domain, we can write the above equation as:

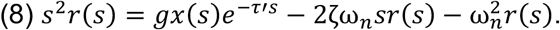

Rearranging the terms gives the ratio of the response to the steering error:

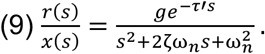

Equation (9) can be interpreted as the transfer function of the system, i.e., the impulse response in the frequency domain. Transforming back into the time domain, the impulse response function is:

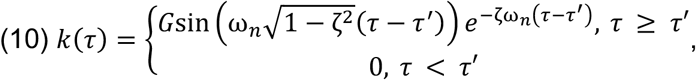

and where

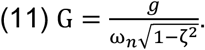

We fit equation (10) to the steering behavior using the MATLAB routine lsqcurvefit to find the set of parameters, ω_*n*_, ζ, *g*, and *τ*′, that minimize the squared differences between the model and observed responses. To assess the significance of the changes in these parameters between experimental contexts, we trained the model on a subset of 200 trials selected uniformly, with replacement, from the data set. We repeated the process 100 times to determine a bootstrap distribution of each parameter value. Outlying fits with any parameter value greater than 2.5 standard deviations from the mean were discarded; no more than 7% of fits were identified as outliers. We used the resulting distribution to calculate the 95% confidence intervals for each parameter as 1.96 times the distribution’s standard deviation. We assessed the significance of parameter changes with a *t*-test.

## 3. Results

To quantify and model the system controlling steering behavior, we analyzed the motor output of monkeys trained to steer through a virtual environment using a joystick to control the angular velocity of locomotion (Figure 1a,b; see Methods). Monkeys learned to control their trajectory through the virtual world for bouts of steering lasting 15-30 s. We analyzed steering behavior in two different target motion contexts. In one context, the target remained at a fixed location in the virtual environment for several seconds before abruptly stepping to a new location to the right or left of the monkey’s heading at the time of the step (Figure 1c). In the second context, the target randomly drifted through the environment over time (Figure 1d). In both contexts, the monkeys learned to match their heading (blue traces) to the direction of the target (red traces).

Differences between the heading and target result in steering errors (purple arrow and traces; Figures 1b-d). Non-zero error signals elicited steering responses (black traces) in the direction of the error. The result of these steering responses is most easily demonstrated by examining the response to the large, transient error to the right occurring just over 5 s from the beginning of the trial in Figure 1c (upward arrow). Following the error signal, the monkey initiated a right steering response indicated by the upward deflection of the black trace. The steering response controlled the rate of change of the monkey’s heading, resulting in a turn toward the target, and a reduction in the subsequent error amplitude.

### 3.1. Steering response to a drifting target is proportional to steering error

Inspection of the steering error and the responses in the example trials from Figure 1c and d suggested that steering responses were approximately proportional to the error and delayed in time, consistent with our previous analysis of steering responses in the step context (22). We sought to confirm this proportional relationship between steering error and response in the drift context. However, unlike the step context, in which step events allowed us to condition responses on an imposed error signal, in the drift context the steering error evolved continuously and randomly. To overcome this difficulty, we parceled steering errors into discrete bins and used this parcellation to condition analysis of subsequent steering responses. Each time that steering error within a given bin was displayed to the monkey, we found the response at 0, 0.21, 0.42, 0.85, 1.69, and 3.39 s after the error was displayed. Repeating this for each analysis bin, we determined the joint distribution of steering errors and responses lagged over time. The resulting joint distributions revealed steering responses that increased with the error with a peak lag of approximately 0.21 s (Figure 2, contours).

**Figure 2.**
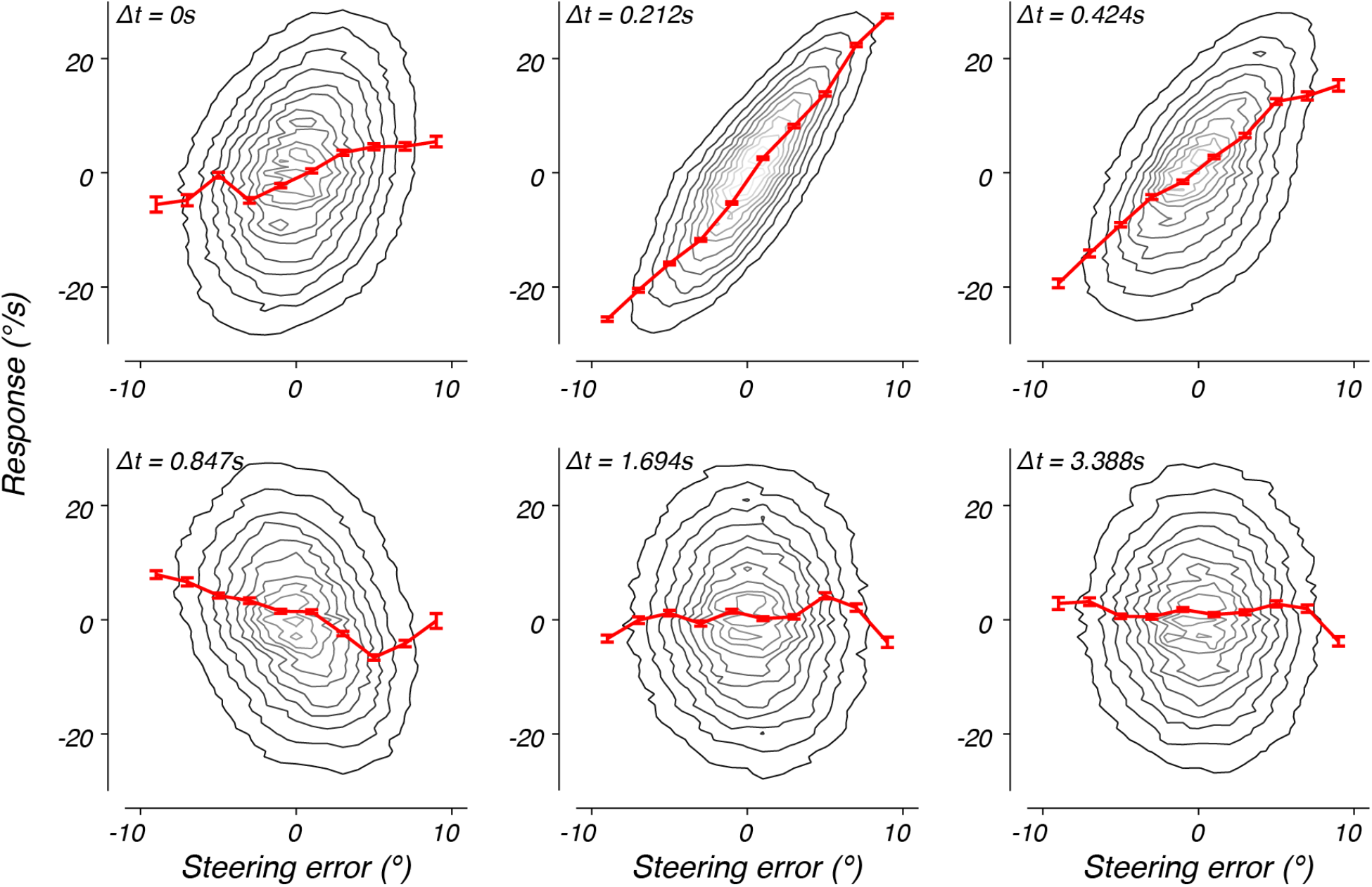
Steering error versus response at several different lags. The joint probability of a steering error and monkey J’s steering response at time lag Δ*t, p*(*x*(*t*), *r*(*t* + Δ*t*)), marginalized across time and trials. The probability distribution is represented by a contour map, with light and dark contours corresponding to higher and lower probability, respectively. The red line plots the mean response, given the steering error was within +/-1 deg of the data point. The error bars represent the standard error of the mean.

By conditioning the responses on a selection of steering errors, we could calculate the mean responses as a function of steering error (Figure 2, red traces). At short lags, the mean response was a nearly linear function of the steering error (red traces). As the lag increased, the shape of the tuning function changed, reversing in sign by 0.85 s. At long lags, the past steering error no longer strongly predicted the response (1.69 s or later for this monkey).

These results suggest that, similar to the step context, the steering response is approximately a linear function of the history of steering errors and evolves dynamically over time. However, the substantial autocorrelation of the steering error over time makes a direct quantification of the steering response function using this method impossible. For example, the distribution at zero time lag (i.e. 0 s) exhibited a weak, but positive correlation to the steering error (Figure 2, top left; monkey F: r = 0.334, p < 0.001; monkey J: r = 0.213, p < 0.001). This relationship arises because the steering error at time 0 is positively correlated with steering errors that occurred just prior to this time point. Therefore, some portion of the observed steering response at this time lag reflects this correlation. To separate the elements of the responses due to the sensorimotor transformation from those that reflect autocorrelations in the stimulus and response, we adopted a nonparametric modeling approach.

### 3.2. Linear model of steering behavior

These observations, combined with previous evidence that steering behavior is feedback dependent (22, 46, 47), suggest an appropriate model of steering behavior is a closed-loop linear model. Several previous investigations of steering behavior have proposed linear feedback models of human steering behavior of the form shown in Figure 3a (23, 25–27). In this class of models, a difference in the target direction, *T*(*t*), and heading, *H*(*t*), leads to an error signal, *x*(*t*). The observed error is sent through a linear response function, *k*(*τ*), which computes the steering response, *r*(*t*), based on a linear combination of the recent history of steering errors. This results in the production of joystick movements that control the rate of change of the monkey’s heading, 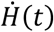 (*t*), proportional to the steering response. The experimental computer integrates these heading changes and the resulting heading signal is once again compared with the target direction, closing the feedback loop.

**Figure 3.**
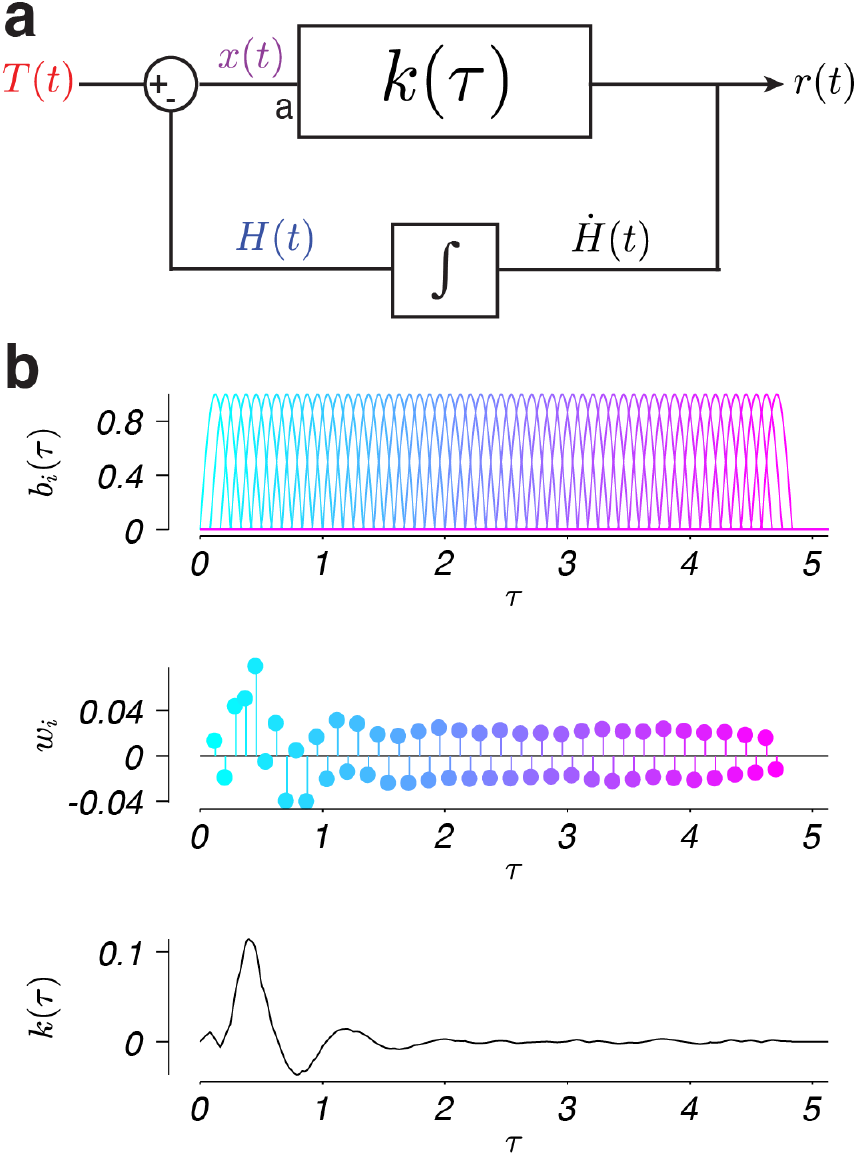
Linear model of steering behavior. a) The monkey observes the current steering error, *x*(*t*), which is the current target position in world coordinates, *T*(*t*), minus the current heading in world coordinates, *H*(*t*). The monkey’s response, *r*(*t*), is modeled as a linear function that sums the weighted past steering errors according to *k*(*τ*). *τ* specifies the temporal delay between the occurrence of a given steering error and the current time, *t*. The response generates a change in heading,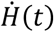, which is integrated over time to generate a new heading. This new heading is then compared with the target position to generate a new steering error, closing the system loop. In the multiplicative noise model we simulated behavior with Gaussian white noise injected at point a. b) The linear kernel, *k*(*τ*), was constructed from overlapping basis functions, *b*_*i*_(*τ*), with *i*indexing functions with peaks at different delays (top; colors). Each basis function was assigned a weight, *w*_*i*_ (middle; colors), and summed to specify a kernel, *k*(*τ*) (bottom). The shown example kernel was fit to data from monkey F in the step context.

Linear feedback models of this form have proven successful at capturing several aspects of human steering behavior. However, these models have undesirable features that prove to be problematic for investigating the neural basis of feedback control. First, the contribution of the history of errors to the linear transformation, *k*(*τ*), depends on both neural sensorimotor integration and biomechanical factors. For example, a key strategy to mitigate uncertainty due to sensory noise is to integrate sensory inputs over time (48). Indeed, results from experimental psychophysics support the conclusion that sensory estimates rely on integration over time (28, 40, 41, 49–51), and one should expect a substantial contribution of sensory integration to the shape of *k*(*τ*). At the same time, the physical constraints of the motor system, such as stiffness and viscosity, result in past motor responses influencing the current response (17–19).

Therefore, *k*(*τ*) can be expected to reflect both neural sensorimotor integration and the physical properties of the plant. However, the results from currently available steering experiments cannot be used to tease apart the relative contribution of sensorimotor integration and motor constraints to steering responses. It is therefore desirable to use nonparametric models to specify *k*(*τ*) such that the form of the weighting given to past errors does not require an exact formulation of the contribution of sensory or motor processing to the response. Second, the assumption of linearity remains untested in most steering contexts, despite the fact that most models leave substantial variance unexplained. It is therefore desirable to derive nonparametric linear models that identify the linear portion of the response with a minimal number of assumptions, such that systematic responses in the residual behavior unexplained by the linear model can be confidently attributed to nonlinearities in the steering system.

We therefore applied a nonparametric method for identifying the weighted linear combination of the history of errors, or kernel, to the current steering response. The kernel (Figure 3b, bottom) was found using a linear combination of basis functions (Figure 3b, top) with weights (Figure 3b, middle) chosen to minimize the squared error between the model response and the actual steering response data (see the *Steering model* section in the Methods). Importantly, this approach makes no assumptions about the integration of error signals over time or the physical constraints of the motor effectors when identifying the kernel, instead it directly estimates the contributions of the linear, nonlinear, and noise-driven components of the overall sensorimotor transformation governing steering behavior.

#### 3.2.1. A linear model captures steering responses in the drift context

We used this approach to fit steering response data in the context of a target that randomly drifts across the virtual world (see Methods, Figure 1d). The best fitting kernels to the steering responses of both monkeys F and J shared similar characteristics (Figure 4a). In response to a brief error pulse, the kernels predict a large response, starting after approximately 0.10 s, in the direction of error, followed by an oscillatory response that decays to zero by approximately 2 s. For a temporally extended, dynamic error input the steering response equals the sum of the kernel response to a continuous stream of impulses of varying amplitude, one for each moment in time.

**Figure 4.**
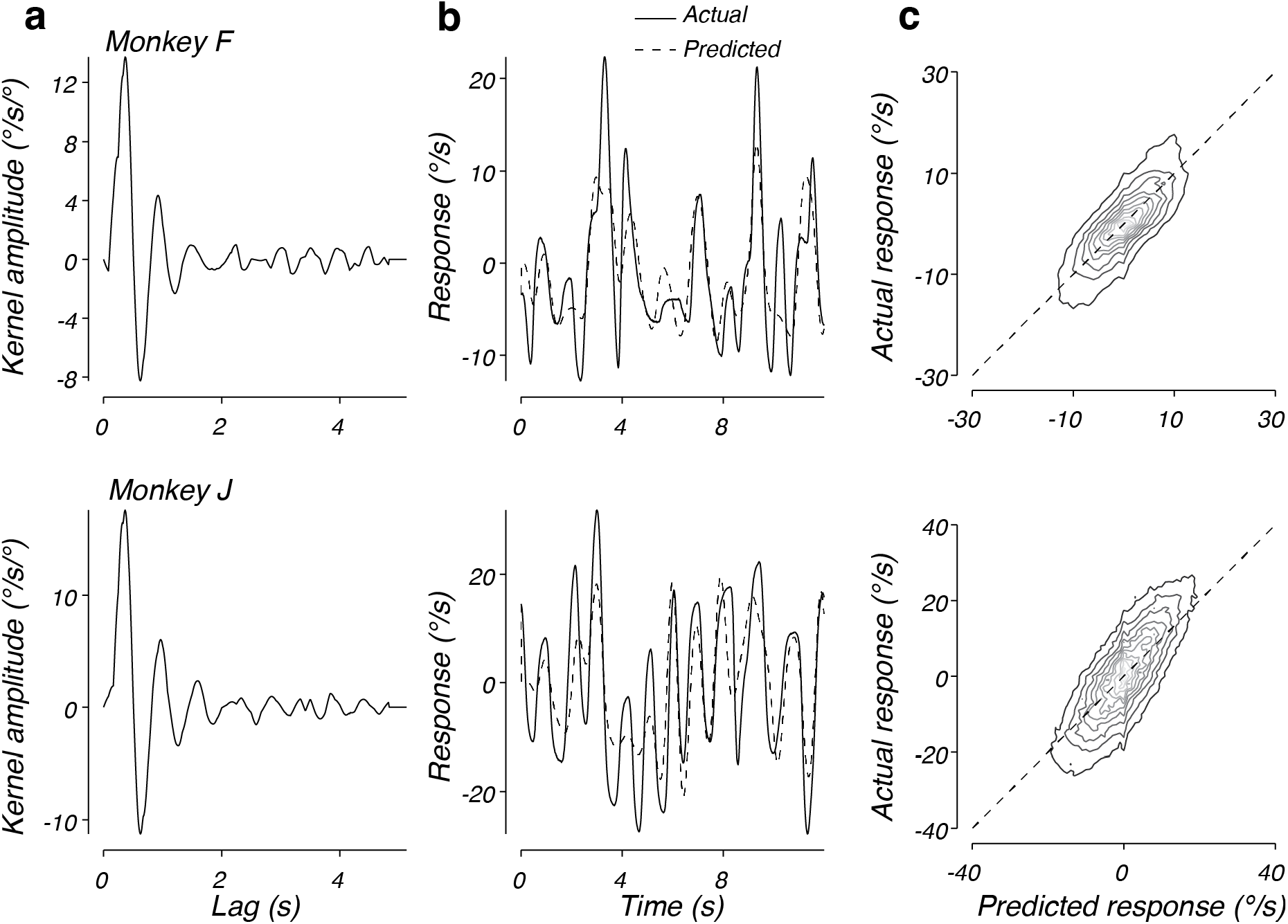
Performance of the linear model on drift data. a) The best fitting linear kernel for monkey F (top) and monkey J (bottom). b) Comparison of the actual and predicted response for 12 s from one trial from monkey F’s data (top) and one trial from monkey J’s data (bottom). Solid lines plot the data from a validation trial and the dotted lines plot the prediction based on the fit to the training data. c) The predicted response plotted against the actual response across time and trials for data from monkey F (top) or monkey J (bottom). Contours plot the joint distribution of predicted and actual responses, with darker lines corresponding to lower probability. Contours are linearly distributed. The dotted line represents unity slope.

Figure 4b plots the actual response (solid lines) and predicted response based on the model (dashed lines) for example trials from monkeys F and J, respectively. In both cases, the model output provided an accurate prediction of the steering behavior. To compare the predictions of the linear model to the observed data across trials, we computed the joint probability of the predicted and actual responses, given the same initial conditions (Figure 4c). The distributions for both monkey F (top) and monkey J (bottom) were aligned along the unity slope line, indicating that the model captured animal behavior well. We quantified this by computing the correlation coefficient between predicted and observed responses and found that the model captured 57% and 56% of the variance in steering responses for monkey F and J, respectively. Interestingly, there were some behavioral responses that tended to be larger than predicted at the extreme response values. These systematic deviations indicate a modest nonlinearity that would be evident when the monkey observes large errors. At smaller response amplitudes, the linear model captured the behavior without systematic errors, but substantial variance remained unexplained, suggesting noise or nonlinearities also contributed to the monkey steering behavior.

#### 3.2.2. Nonlinearities within the step context are small

Teasing apart the contributions of nonlinearities and noise to sensorimotor processing requires an approach that can isolate the systematic components of the steering response from the components of the response that are not systematically related to the steering error. A straightforward method for removing the nonsystematic portions from the steering response is to average steering responses to identical error inputs. Because the steering system is inherently closed-loop, animal behavior contributed significantly to the steering errors and controlling the sequence of error inputs was not possible in the drift context. We therefore turned our analysis to behavior in the step context, which was explicitly designed to control the steering error delivered to the monkeys. In the step context, the target was moved in world coordinates discretely to generate a specific error (e.g. 25 deg) regardless of the steering behavior. This allowed us to determine the mean steering response to specific error input in a small temporal window around each target step (Figure 5) (22).

**Figure 5.**
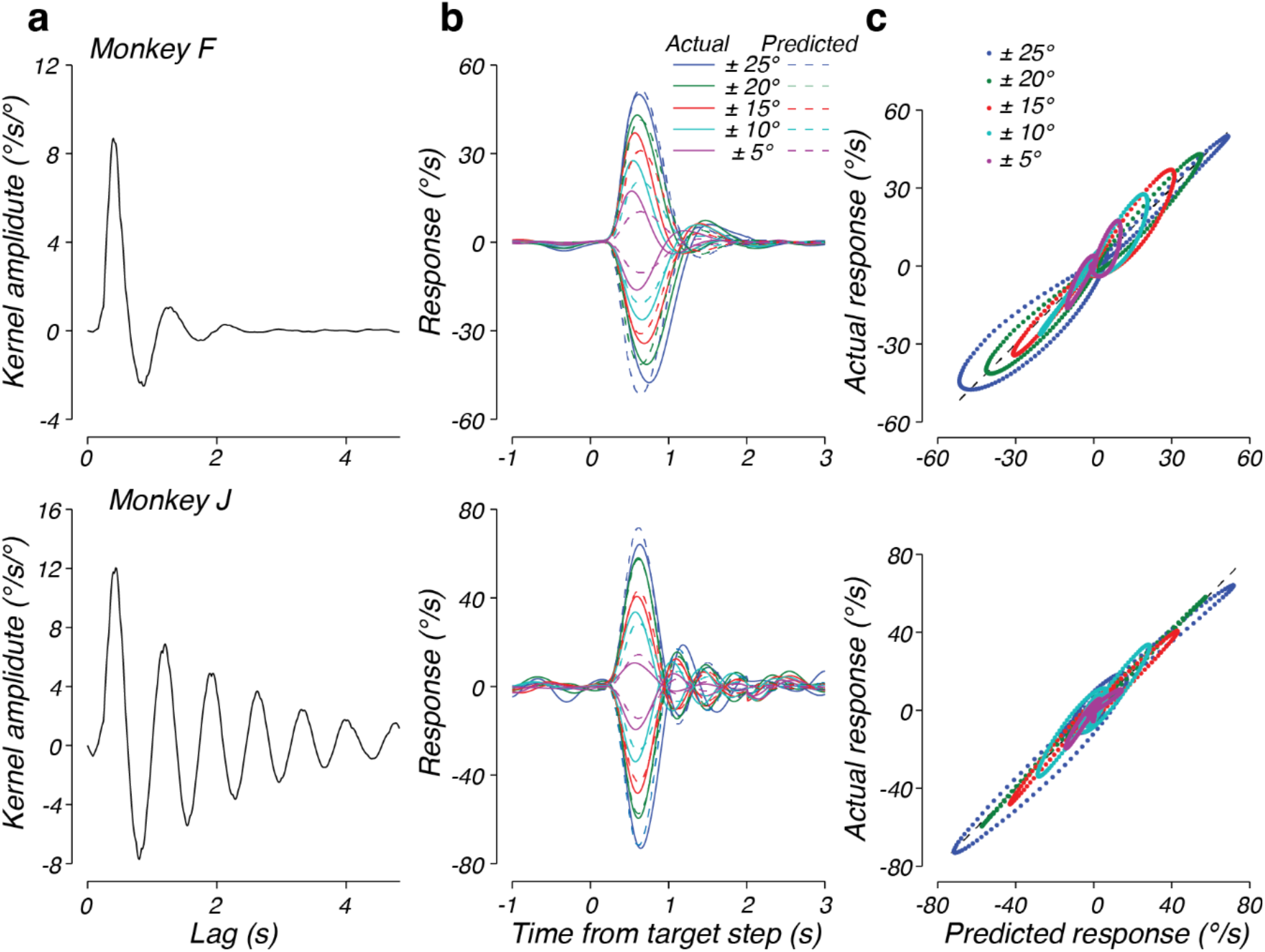
Results of the linear model fit to trial-averaged data from the step context. a) The best fitting linear kernels for monkey F (top) and monkey J (bottom). b) Comparison of the mean response to steps of different amplitudes to the predicted response over time. The solid lines represent the actual data, the dotted lines plot the prediction. The color of each trace represents the amplitude of the target step from which the data were averaged. Results for monkeys F and J are presented in the top and bottom panels, respectively. c) The predicted response plotted against the actual response. Different colors refer to different amplitude target steps.

We leveraged the mean steering responses to target steps to evaluate the capacity of the linear model to capture the systematic portions of steering behavior. Importantly, a linear system will respond to error inputs of different amplitudes with identical, scaled responses. Therefore, a perfectly linear system should capture the mean steering response across step sizes. The best-fitting kernel for each monkey is shown in Figure 5a. Similar to the kernels identified in the drift context, the kernels fit to the mean step responses were characterized by a large onset response followed by a damped oscillation.

Direct comparison of the responses of the linear model to the mean step responses revealed that, overall, a linear model fit the behavior quite well (Figure 5b). Across step sizes, the identified kernel captured 93% and 94% of the variance in the mean responses for monkeys F and J, respectively. Close examination of the linear model output (dashed lines) and actual responses (solid lines) revealed that the remaining unexplained variance results from small deviations between the linear prediction and behavior. Plotting the predicted response against the actual response revealed that the deviation from the linear prediction came in three forms (Figure 5c). First, there were systematic differences between the residuals to left (negative responses) and right (positive responses) steering errors, perhaps resulting from the asymmetries in the muscle groups of the wrist (52). This indicates a nonlinear interaction between steering error and response direction. Second, the linear model systematically underestimates the peak amplitude of responses to small target steps, indicating a modest nonlinearity in the amplitude of the response to a target step. Third, unlike the model, the timing of the peak responses in the data tended to be earlier for small steps and later for large steps. These two latter deviations may be a signature of un-modeled compensation for a component of noise that increases with signal size (53)(see Discussion).

While the linear model successfully described behavior in both the step and drift context, we observed substantial differences between the kernels identified between the two contexts. For example, when one compares the kernel identified from monkey F in the drift context (Figure 4a, top) and the kernel identified from monkey F in the step context (Figure 5a, top), the oscillation observed in the kernel fit to monkey F’s behavior was slower in frequency and decreased in amplitude in the step context, with a similar time constant of decay of the envelope of the oscillation. Monkey J also had slower frequency and decreased amplitude oscillations in the step context, but had a much slower time constant of decay of the envelope of the oscillations (compare Figure 5a, bottom and Figure 4a, bottom). These results suggest that the response function deployed by each monkey depended strongly on the experimental context (see below).

Because we applied our identification analysis to trial-by-trial data from the drift context but trial-averaged data for the steps, the observation that the kernels changed between the two experimental conditions could be an artifact of the difference in analysis. Therefore, we verified that the kernels identified from the mean step response data were robust to our analysis method. To do so, we fit the kernels to the trial-by-trial responses in the step context, as in the drift context. The resulting kernels matched the kernels found by fitting the mean step responses (Figure 6a), confirming that the kernel differences were not due to our analysis technique. However, examination of the trial-by-trial predictions revealed that the ability of the linear model to capture the corresponding behavioral data differed between monkeys. The linear model performed well for monkey F on individual trials (c.f. Figure 6b, top), capturing 70% of the variance in the monkeys responses (Figure 6c, top: *r*^*2*^ = 0.70). In contrast, inspection of individual trials from monkey J revealed that the actual response deviated significantly from the response predicted by the linear model (Figure 6b, bottom). In particular, large oscillations often occurred in the behavior when the linear model predicted little or no response. Plotting the response prediction versus the actual response reveals that the largest responses often occur when the linear model predicts little or no response (Figure 6c, bottom). Across trials and time points, the linear model captured only 18% of the variance in this monkey’s steering responses. Because the model captures the mean response to steps of the target as well as for the other monkey, our inability to predict trial-by-trial behavior suggests a substantial contribution from noise (or possibly a complex nonlinearity) that reverberates through the steering system on a trial-by-trial basis.

**Figure 6.**
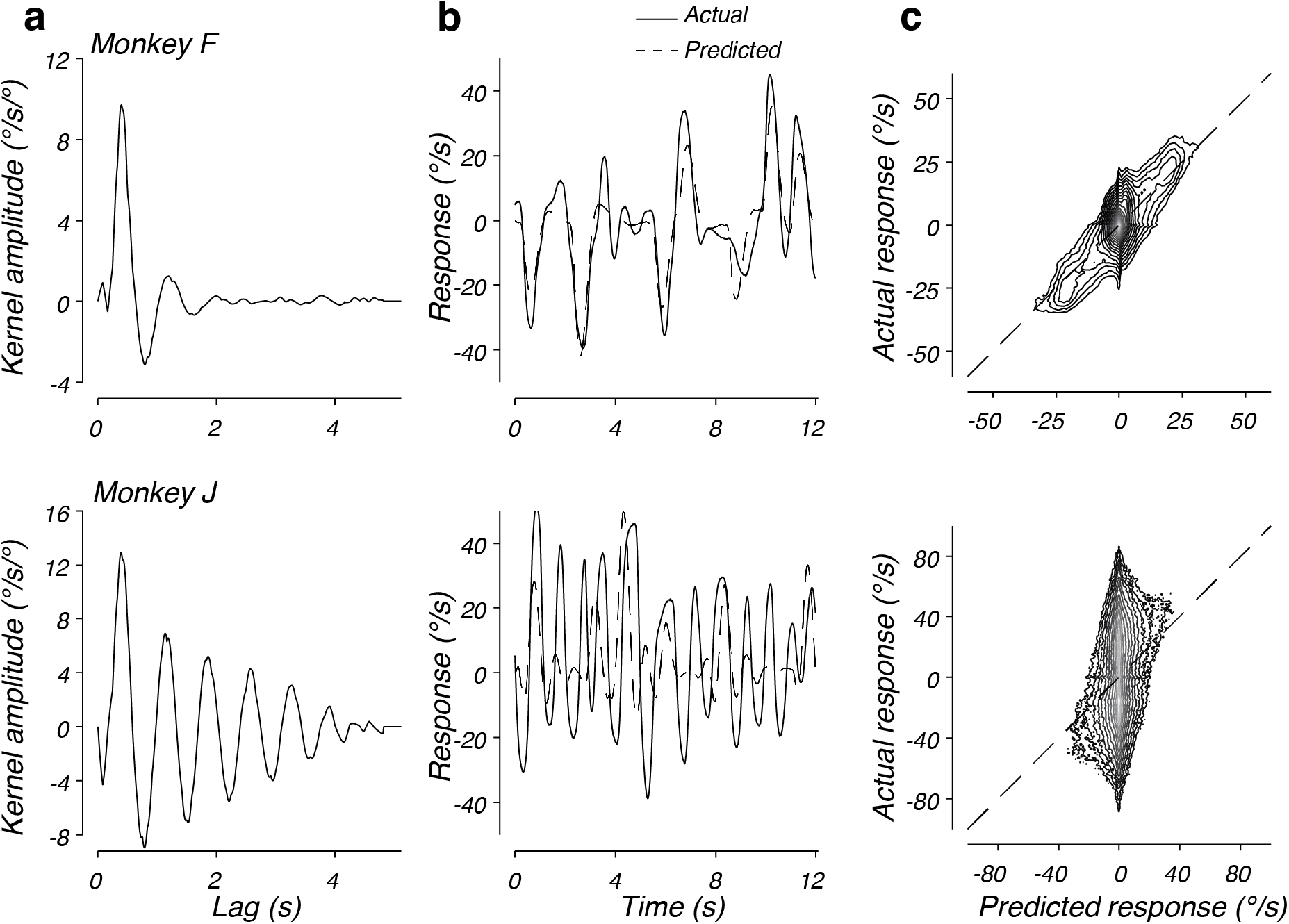
Performance of the linear model on step data. a) The best fitting linear kernel fitted to trial-by-trial steering responses during the step context for monkey F (top) and monkey J (bottom). b) Comparison of the actual and predicted response for 12 s of one trial of monkey F’s data (top) and one trial of monkey J’s data (bottom). The solid lines represent the actual data; the dotted lines plot the prediction. c) The predicted response plotted against the actual response. Conventions as in Figure 4. Contours are logarithmically distributed.

Taken together, the above analyses confirmed previous results suggesting that a linear model can capture the step-averaged portion of the steering response (25–27, 29). However, the identified kernels differed substantially from those found for the drift context, indicating that experimental context strongly impacts the shape of the response function. At the trial-averaged level, the linear models capture well over 90% of the variance of the step behavior, with the remaining unexplained variance largely associated with modest nonlinearities in the response kinematics to left and right steering responses and steps of different amplitudes. At the single-trial level, substantial variability was observed (30-82% unexplained variance) that broadly resembled the level of unexplained variance observed in the drift context (43-44% unexplained).

### 3.3. Residuals analysis

We next sought to determine the sources of variation left unexplained by the modest nonlinearities observed within a behavioral context. Because our analysis relies only on the initial condition of the steering response to predict all subsequent behavior, a substantial proportion of the unexplained variance likely reflects the accumulation of errors in prediction due to factors that are not systematically related to the steering error. Therefore, we analyzed the residual steering behavior not explained by the linear model for each monkey and task condition.

Comparison of the actual steering responses and those predicted by the linear model on individual trials indicates that residual responses have temporal correlations that extend over several hundred milliseconds (e.g. Figure 6b, bottom), suggesting the presence of a noise process that, after reverberating through the steering system, gives rise to substantial low frequency residuals. To quantify this, we computed the frequency content of the residual steering responses for each monkey in each behavioral context. Despite the large differences in the total unexplained variance, normalizing the residual spectra by their total power revealed shared characteristics across monkeys and contexts (Figure 7a, black lines). Residuals in all cases had substantial power at low frequencies, gradually increasing up to a peak at approximately 1 Hz. Power in frequencies higher than 1 Hz quickly decreased, becoming small at frequencies larger than 3 Hz. These results suggest that the unexplained variance in responses arises from a source of stochasticity that is similar across monkeys.

**Figure 7.**
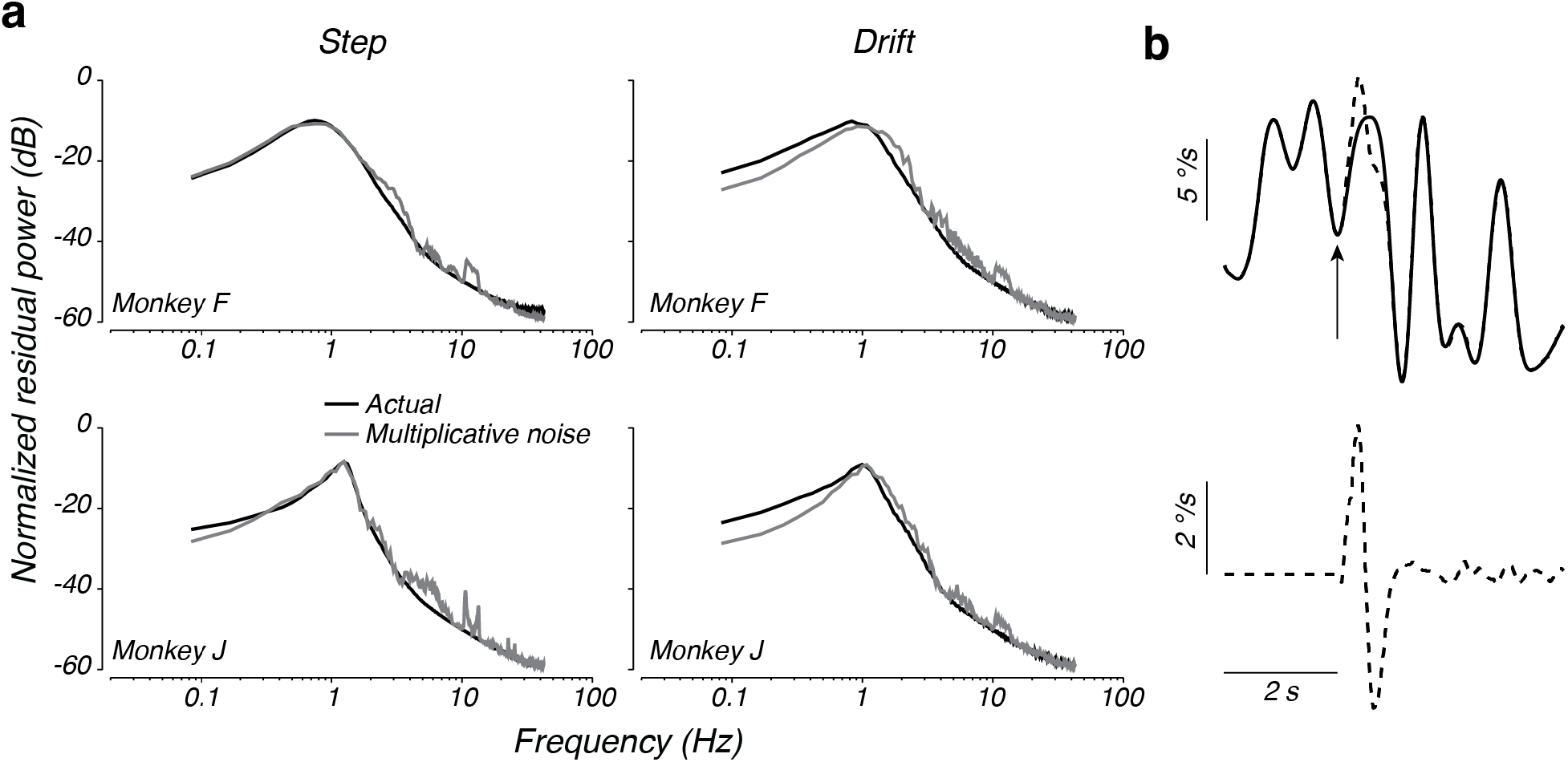
Residual spectra for each monkey and context. a) Spectra of the residual behavior not explained by the best-fitting linear model (black), spectra of the residuals of the multiplicative noise model (gray). Each spectrum is normalized by its total power. b) *Top*: Simulated response using the kernel found for monkey F in the drift context (solid trace) and the response to an identical trial, except for a 20 deg perturbation of the steering error applied at the time of the arrow (dashed line). *Bottom*: the isolated response to the perturbation, obtained by subtracting the simulated response without the perturbation from the simulated response with the perturbation.

Given that the systematic portion of the monkeys’ steering behavior was well explained by a linear model, we hypothesized that the form of the spectra of the residuals could be explained by considering the effect of the kernel operating in a closed loop on sensorimotor noise. In a linear feedback system, the response of the system to noise reflects not only the feedforward kernel, *k*(*τ*), but also computations in the feedback loop. In the case of steering, the feedback loop performs integration of the motor output (Figure 3a). Thus, we expect the spectra of the residuals to be shaped by the closed-loop transfer function, even for broad spectrum noise. Consideration of the effect of the closed-loop transfer system on noise analytically confirms this intuition (see Appendix), but to illustrate the effect, we used our nonparametric kernel fits to computationally generate the expected response of the system to a drifting target (Figure 7b, top, solid line) as well as an identical drifting target plus a brief pulse (introduced at the time of the upward arrow in Figure 7b) to simulate the effect of a noise perturbation on behavior (Figure 7b, top, dashed line). The difference in the response with and without the pulse reflects the closed-loop impulse response to a noise perturbation (Figure 7b, bottom; alternatively, the impulse response could be derived analytically from *k*(*τ*), Appendix, eq. 16). The resulting impulse response exhibits an oscillation with a dominant frequency close to 1 Hz, much like the empirical residual spectra. This simulation illustrates how the peak in the spectra of the residuals in Figure 7a could arise even for flat or broadband spectral noise.

We formalized this analysis by considering a model in which the error observed by the monkey is subject to noise before convolution (multiplication in the frequency domain) by the steering kernel (Methods, Eq. 1; Figure 3a, location a). We tested how this ‘multiplicative’ noise model impacts the spectrum of residuals by simulating steering behavior using kernels fit to the behavior, with Gaussian white noise added to the steering error. Following the model simulations, we determined the spectrum of residuals by subtracting the simulated responses with noise from the response predicted by the kernel alone.

Comparison of the spectra of the residuals from the multiplicative noise model and the observed residual power revealed a striking similarity across monkeys and experiments (Figure 7a, gray lines). The multiplicative noise model correctly predicted the residual power to increase with frequency up to a peak at about 1 Hz, with residual power falling off dramatically after this peak. This result was confirmed by an analytic treatment of the closed-loop steering system, which demonstrates how the spectrum of the noise is shaped by the power spectrum of the kernel (Appendix, Eq. 19). This analysis demonstrates that a multiplicative noise model accounted for most of the response variance left unexplained by the linear model.

Although the multiplicative noise model captured the general shape of the residual spectra, it mildly underestimated the power of the residuals at low frequencies and overestimated the power at high frequencies. We therefore sought to more directly test the multiplicative noise model, by inferring the spectrum of noise directly from the data. Assuming a linear model with multiplicative noise that is uncorrelated with the location of the target in world coordinates, the spectrum of the noise can be estimated from the residuals, linear response, and input spectra (45) (see Appendix). The inferred noise spectra across monkeys and contexts were highly similar after normalizing each by the total variance of the steering error (Figure 8, small closed and open circles). In all cases, the estimated noise spectra were approximately white in the middle range of frequencies (∼0.3-2 Hz), consistent with our simple multiplicative noise model. However, the estimated noise spectra in lower or higher frequency bands decreased with frequency, suggesting that a model that assumes white noise added to the steering error before filtering by the steering system misses some characteristics of the noise process within the sensorimotor systems responsible for steering.

**Figure 8.**
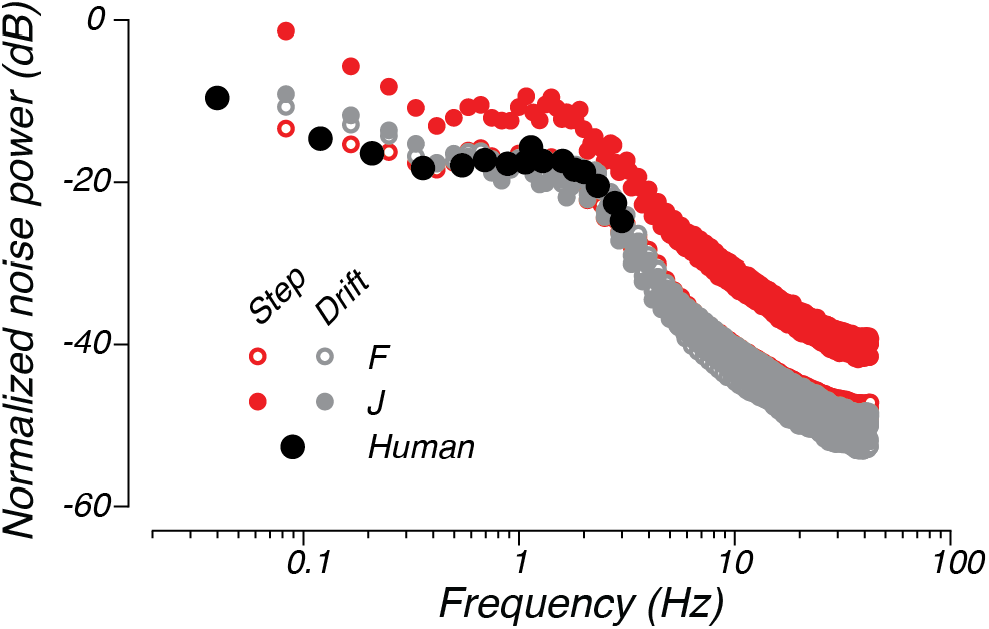
Estimated spectra of multiplicative noise. The small open circles and small closed circles plot the multiplicative noise spectra estimated for monkeys F (open circles) and J (closed circles), respectively. The red points plot the estimate from the step context and the gray points plot the estimate from the drift context. The large black circles re-plot the data from Jex and Magdaleno (54) for a human manual control task with input taken from a continuous spectrum, much like the input for our experiments. Each spectrum is normalized by the error variance in the corresponding task.

While the total power of the noise differed across monkeys and contexts, the noise spectra were nearly identical for monkeys F and J when normalized by the variance in the error signal observed by the monkey. This similarity suggests that the nondeterministic steering responses observed across monkeys and experiments results primarily from a source of noise that scales with the error variance. Only the noise spectra estimated from monkey J in the step context differed from the other normalized spectra. The increase in the noise fraction for monkey J in the step context suggests that another source of stochasticity that is independent of error variance contributed to the steering responses for this monkey and context.

Interestingly, the normalized multiplicative noise spectrum was extremely similar to that estimated for humans performing manual tracking tasks. The large black dots in Figure 8 re-plot the normalized noise spectrum for data taken from human subjects performing a manual control task (54). The obvious similarity between the human manual control and monkey steering data suggests that variance in steering behavior stems from similar sources of noise across animals.

Taken together, our analysis of the residuals has strong implications for the origins of noise during sensorimotor function. Our multiplicative noise model is consistent with noise occurring in the early processing stages of the steering system, consistent with previous conclusions that variance in sensorimotor behavior originates mainly during sensory processing (43, 44).

### 3.4. Experimental context induced nonlinear transitions in control policy

Across experiments and monkeys, the identified kernel took the form of a damped oscillation. However, comparison of the kernels fit to each monkey revealed changes in the features that characterize this damped oscillation between contexts (compare Figures 4 and 6). These changes appear to affect more than the gain alone, indicating nonlinearities in the sensorimotor transformation with respect to context. To evaluate this further, we leveraged the fact that a kernel in the form of a damped oscillation matches models of manual control in which the acceleration of the hand that controls the steering response is proportional to a linear combination of the steering error, a spring-like restoring force parameterized by the spring’s stiffness, and a viscous damping term (23, 29, 55–57). Therefore, the parameters of a fit of the steering responses in each experimental context to such a second-order linear model allow for a straightforward interpretation of how the kernel changes between contexts.

We validated the second-order linear model using two analyses. First, we fit the model to the raw data. Then we transformed the fit second-order model into a kernel response function (see *Second-order linear model* in Methods) to directly compare the associated kernel with those identified using our nonparametric model (Figure 9a). Across both monkeys and contexts, the second-order linear model and kernel fit to the behavior exhibited very similar dynamics – the impulse response of the second-order linear model captured 90-99% of the variance in the regression kernel impulse responses. Modest differences in the delay and onset kinetics account for the majority of this difference. Next, we evaluated the overall performance of the second-order and nonparametric kernels in predicting monkey behavior. The second-order model explained nearly exactly the same fraction of variance in steering responses as the kernels identified by the nonparametric approach (Figure 9b). This strong agreement between the kernels identified using our nonparametric methods and the second-order models that have previously been used to explain steering behaviors provides compelling evidence that the second-order model accurately describes monkey steering behaviors.

**Figure 9.**
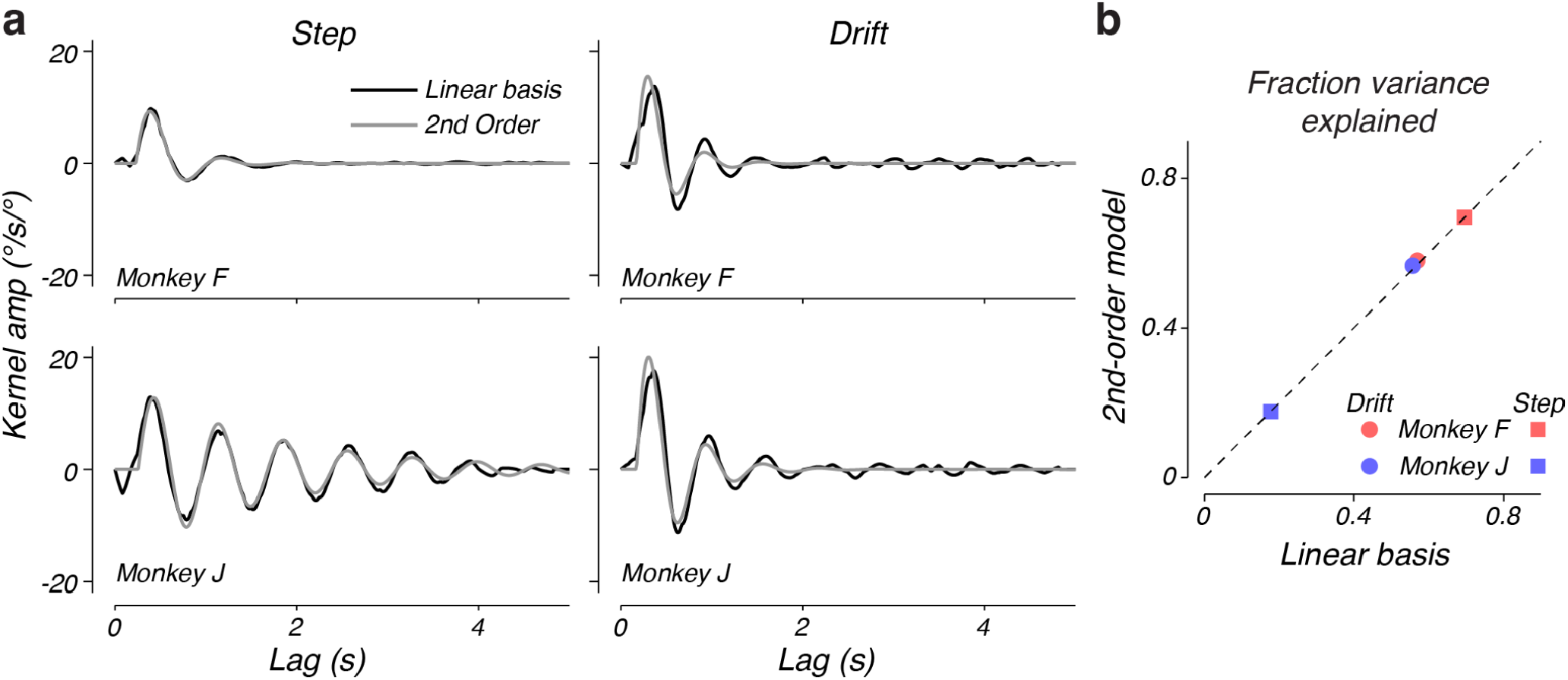
A second-order steering model is consistent with identified kernels. a) Comparison of the kernel, *k*(*τ*), found by the nonparametric linear basis and the kernel corresponding to the best-fit second-order linear model. Left and right panels indicate the fits to the step and drift contexts, respectively. b) Comparison of the fraction of behavioral variance explained by the nonparametric linear basis model and the best-fitting second-order linear model for each monkey and experiment.

Having validated the second-order model as a parametric description of the steering response, we next compared the values of the parameters fit to each experimental context for each monkey. As expected, based on visual inspection of the kernels, the parameters of the linear system of each monkey changed with experimental context (Table 1). To assess the significance of these changes, we performed a bootstrap analysis of fits to the data. For almost all parameters in both monkeys, the differences were highly significant (Table 1), demonstrating the changes observed in our kernels between contexts were not due to chance. Notably, we did not only observe changes in the gain and time delay, but we also observed significant changes in the damping ratio and natural frequency parameters ζ and ω_*n*_, parameters typically considered to be static properties of the motor effectors. For monkey J, the damping ratio differed most between contexts, decreasing by ∼69% from the drift to step context. In contrast, the damping ratio for monkey F did not change significantly in the step context relative to the drift context. For both monkeys the undamped natural frequency significantly differed between contexts, decreasing 14 and 20% from drifts to steps for monkeys J and F, respectively.

**Table 1.** Changes in the steering system of the monkeys quantified by the fits to a second-order linear model. Values of the damping ratio (ζ), undamped natural frequency (ω_*n*_), gain (*G*), and time delay (*τ*′) indicate the mean +/- 95% confidence intervals, which were calculated from the standard deviation of the bootstrap distribution. The percent change is calculated relative to the drift context. Significance was assessed by a two-tailed *t*-test based on the ratio of the change in mean parameter values to the square root of the summed variances of the bootstrap distributions of each parameter value (see Methods).

| | | $\zeta$ | $w_n$ (rad/s) | $G$ | $\tau'$ (s) |
| --- | --- | --- | --- | --- | --- |
| F | Step | 0.34 +/- 0.02 | 8.51 +/- 0.24 | 0.18 +/- 0.01 | 0.23 +/- 0.01 |
|  | Drift | 0.31 +/- 0.04 | 10.67 +/- 0.53 | 0.29 +/- 0.02 | 0.17 +/- 0.01 |
|  | Percent change | 8.72 | -20.24 | -38.01 | 35.42 |
|  | t(99) | -1.32 | 7.26 | 11.03 | -8.03 |
|  | p | 0.190 | <0.001 | <0.001 | <0.001 |
| J | Step | 0.07 +/- 0.02 | 8.86 +/- 0.19 | 0.17 +/- 0.01 | 0.26 +/- 0.01 |
|  | Drift | 0.23 +/- 0.03 | 10.33 +/- 0.35 | 0.33 +/- 0.02 | 0.17 +/- 0.01 |
|  | Percent change | -69.49 | -14.21 | -49.79 | 48.92 |
|  | t(99) | 8.55 | 7.16 | 16.02 | -10.34 |
|  | p | <0.001 | <0.001 | <0.001 | <0.001 |

## Discussion

Because system output influences subsequent input, identification of sensorimotor transformations in a closed-loop context has proven a difficult challenge in neuroscience (42, 58). This challenge has significantly limited progress in investigating the neural mechanisms implementing control policies in the natural context of closed-loop control. Here, we overcame this challenge by adapting nonparametric, kernel-based approaches to model the sensorimotor transformations in a closed-loop behavior directly. Using this approach, we were able to accurately identify the linear transformation of sensory input into a motor output, without being limited by a chosen model of sensory integration or motor constraints.

We focused our analysis on steering responses exhibited by monkeys trained to track targets, a sensorimotor system which relies heavily on sensory feedback to guide behavior in a closed-loop (22, 46). Using our nonparametric approach, we found that a systematic linear transformation could explain between 18 and 70% of the variance in the observed steering responses. The form of this linear transformation was consistent with previous modeling efforts that assume steering behavior can be approximated as an instantaneous gain on the time-delayed steering error constrained by viscous and spring-like forces (23, 25, 26, 29). However, our analysis revealed two features of the transformation of steering error by the nervous system that suggest a more complex interpretation. First, there was surprisingly large trial-by-trial variability that, despite its large amplitude, could be accounted for by a simple model of noise in the neural representation of steering errors. Second, we observed significant changes in the parameters associated with viscosity and stiffness with experimental context, two features that are normally ascribed to the physical properties of the shoulder, arm, and wrist. These results suggest a more general interpretation of viscosity as a resistance to changes in the steering response and stiffness as a resistance to non-zero steering output. Notably, instead of reflecting properties of the motor effectors, these features may reflect flexible processing by the neural systems underlying the integration of sensory information and its transformation into a set of motor commands.

### 4.1. A linear feedback system supports steering responses

Several different models have been proposed to describe steering behavior in humans and other animals. Our model assumes a dynamic, linear feedback system similar to other studies that have successfully modeled steering and other navigation behaviors (23, 25–27). However, it should be noted that other models have proposed simple heuristics that are used to guide navigation. For example, navigation to a goal can be achieved by setting a curved path to the goal and maintaining that path by keeping the retinal velocity of the target constant (59). Other approaches include controlling the time to zero steering error such that it equals the time of collision with an object (60, 61) or integrating flow information over time to measure one’s path (59, 62). However, each model makes distinct predictions and, when compared directly to actual steering behavior, a linear feedback system was previously found to best describe steering behavior (23).

The exact form of the dynamic linear system used to model the steering system often differs from experiment to experiment. These differences can be quite subtle and the differences between their predictions can be difficult to quantify. Our kernel-based method largely supported a relatively simple model with steering responses driven by a time-delayed steering error, a viscous-drag like resistance to the hand motion driving steering responses, and a spring-like restoring force acting on hand position. However, there are at least two ways that central neural processing likely contributes to the shape of the kernel that cannot be captured by such a simple model. First, related steering tasks have found a strong dependence of steering on the reliability of sensory information (21, 26, 51). Similar effects have been documented in other manual control tasks (40, 41) and smooth pursuit behavior (63, 64) and are generally consistent with the principle of Bayesian integration in sensorimotor behavior (65, 66). Therefore, the simple second-order model likely requires augmentation to account for the effects of stimulus reliability on the sensorimotor system governing steering. Second, the physical properties of the motor system are adjusted according to the context under which a behavior is being executed (31), suggesting flexible changes in motor policies must also be incorporated into models of steering behavior. Our approach provides a simple method for quantifying changes in sensorimotor transformations due to the effects of stimulus reliability or context and can form a basis for comparison of the predicted transformation functions of models proposed to capture these effects (67).

### 4.2. Residual steering responses suggest a sensory source of noise

We used a nonparametric approach to identify the linear portion of the steering system. For a given context, this linear description fit the systematic steering responses very well. This enabled us to use a simple linear model to estimate the temporal statistics of the residual responses. The resulting residual behavior was found to be well fit by multiplicative noise that is shaped by the closed-loop response, resulting in a peak in residual power near 1 Hz. This was largely consistent with previous models of manual control (45), and matched human variability at a striking level (54).

The most straightforward interpretation of multiplicative noise in our model is variability arising in the measurement of the steering error by the visual system. Our results therefore suggest that the majority of noise in the steering system can be attributed to sensory noise, consistent with the conclusions from the analysis of smooth pursuit eye movements, where much of movement variability can be attributed to variability in the encoding of speed, direction, and timing of visual input (43). In smooth pursuit, this variability is tightly linked to noise in the encoding of motion in the middle temporal cortex (68) that is transmitted to downstream neural populations (69). Given our results linking the sensitivity of individual sensory neurons in the medial superior temporal (MST) area of the cortex to that of steering behavior (70), this interpretation predicts that the residual activity of individual neurons in MST should (1) correlate with steering responses and (2) have noise statistics that are approximately white and uncorrelated over time, up to a high frequency cutoff.

However, the conclusion that steering variance arises from noise in sensory encoding should be tempered against two other general interpretations that we cannot rule out with our current data. First, previous work has suggested sources for variability in sensorimotor systems that are not sensory in origin. Analysis of neural activity in the interval between sensory input and motor execution has revealed substantial response variability (14, 71). Indeed, higher order systems such as those responsible for planning a motor response (72) or setting the strength of sensorimotor transformations (73–75) have recently been related to behavioral variation. Further, it is important not to discount the possibility of noise in motor execution as a substantial source of variability in other sensorimotor behaviors (53, 76–81). Second, complex nonlinearities not modeled by a linear system might contribute significantly to variance unexplained by the linear model. For example, a nonlinear interaction between limb state and steering error would amplify variance that originates as sensory noise and might explain poor model fits for monkey J in the step context.

### 4.3. Nonlinear contributions to steering behavior

Our ability to accurately determine the linear contribution to the steering response also allowed us, for trial-averaged data, to quantify nonlinear contributions to steering. This analysis revealed that, within a given steering context, the contribution of such systematic nonlinearities made up no more than 10% of the total response variance. While trial-averaging was only possible in the step context, we expect similar contributions of nonlinearity within the context of the drift context based on the results of our residuals analysis in those experiments. Some of this nonlinear response could be attributed to the systematic asymmetries in the leftward and rightward steering responses of the monkeys, which likely result from biophysical constraints such as asymmetries in the pulling directions of different muscle groups in the wrist (52). However, there were also small deviations in the exact response kinematics from the linear system, even when considering only one steering direction. Close examination of the residual behavior revealed that the linear model tended to underestimate the peak amplitude of responses to small targets and missed the systematic trend for larger steps to have delayed peaks. These differences could either reflect the impact of nonlinearities on the initiation of steering responses or could reflect a dynamic change in motor policy reflecting optimization over time (82, 83), similar to that observed in the saccadic eye movement system and attributed to the mitigation of signal-dependent motor noise (53). Future modeling and experimental efforts directed at understanding the interaction of biomechanical constraints and motor optimization are required to tease apart the contribution of each to the steering response.

While the contribution of nonlinearities to steering responses within an experimental context was small, our nonparametric method revealed more dramatic nonlinearities across experimental contexts. Quantifying these differences by fitting a second-order linear system to the data, we found that context impacted the gain and time delay of steering errors, consistent with results from previous steering (21, 26, 29) and manual control (55, 84, 85) experiments. However, we also observed significant changes to terms associated with the parameters representing viscous drag and spring-like forces of the motor system. These parameters are typically assumed to reflect biomechanical properties of the arm, leading previous work to assume these parameters remain static across experimental contexts (23, 26, 29, 57, 86, 87). One possible interpretation of our results is that the resonant frequency and level of damping of the arm change given task instructions, as has been observed in previous experiments directed at determining the biomechanical properties of the wrist (31). However, given that even reflexive behaviors are flexibly adapted to the current context by central mechanisms (88), it is also possible that these classically biomechanically interpreted parameters additionally reflect central neural processing that may change as an animal adapts its sensorimotor transformations to the current behavioral context (89). Future experiments that record EMG signals during steering behavior will be helpful in teasing apart these hypotheses.

While our approach allows us to determine how closely a linear model can explain the steering responses, as formulated it cannot help to identify the nonlinear components of the system without making further assumptions. Formally, one can use a Volterra series expansion to determine second-, third-, and higher-order contributions to the behavior, but such expansions require amounts of data that increase rapidly with the degree of the higher order terms (90). Alternatively, one can make simplifying assumptions about the nature of the nonlinearity, such as a static nonlinear transformation following the linear transformation (35). Indeed, static nonlinearities have been leveraged to improve model fits in similar sensorimotor behaviors (28, 30, 32, 91).

### 4.4. General applications to sensorimotor neuroscience

We expect that the nonparametric approach presented here and in other recent work (40–42) will be useful to other applications where sensorimotor behavior must be studied within a closed loop. A key challenge in the study of closed-loop behaviors is that experimenters often must make assumptions as to the form of the model that will explain the systematic and nonsystematic components of that behavior. Such assumptions include the use of Bayesian integration (51), Kalman filters (28, 40, 49, 92), and optimal feedback control laws (82, 93, 94). These can be highly powerful in providing normative explanations for a wide range of behaviors, but caution must be exhibited as their assumptions can be difficult to validate (95). Adopting a nonparametric approach can act as a complement to model-based analysis by helping to guide the selection of possible models for comparison and the experiments that will best differentiate between them.

Finally, we anticipate that our nonparametric method will prove to be valuable to the study of the neural mechanisms underlying closed-loop sensorimotor control. Models built at the level of behavior have proven extremely useful for understanding the computations and algorithms used by the brain to implement sensorimotor control. However, because neural systems rely on several, nonlinear processing stages (96–98) and represent important behavioral variables in mixed and dynamic populations (7, 99, 100), it is often difficult to translate the activity of neurons and networks to concepts developed at the level of behavior (101). We propose that developing kernel functions of sensorimotor behavior will help bridge that gap by providing a common language to the study of behavior and the study of neural responses (70).

## Appendix

*Approximation of the spectrum of multiplicative noise*. Under a linear model with multiplicative noise, the steering response can be specified in the Laplace domain as:

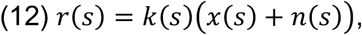

where *s* is a complex number and corresponds to the frequency parameter. *x*(*s*) is the steering error, defined as *T*(*s*) −*H*(*s*) (i.e., the target direction minus the heading), *n*(*s*) is the noise input, and *k*(*s*) is the linear gain as a function of frequency. The heading is defined as the integral of the response, which in the Laplace domain can be written as

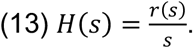

Inserting equation (12) into equation (13) and rearranging, we can parcel the response into the closed-loop response to the target and the noise

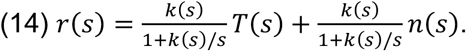

Therefore, the deterministic portion of the response, 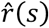, is defined as

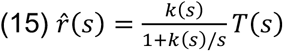

and

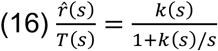

is the closed-loop kernel of the system. In this case, the power spectrum of the deterministic response can be defined as

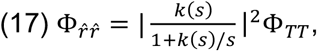

where Φ_*TT*_ is the power spectrum of the target.

The residual behavior due to noise will be

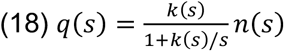

and the power spectrum of the residuals, Φ_*qq*_, is defined as

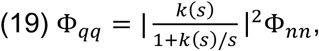

where Φ_*nn*_ is the power spectrum of the noise input. Combining equations (17) and (19), we can define the power spectrum of the noise input as

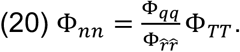

Therefore, to estimate the noise spectrum under the multiplicative model, one can combine the measured power spectra of the input, Φ_*TT*_, the estimated power spectrum of the deterministic response from the response expected based on the kernel, 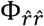, and the measured power spectrum of the measured residuals, Φ_*qq*_.

## Data availability

Data and accompanying data analysis routines are available from the corresponding author upon reasonable request.

## Acknowledgements

We thank Daniel J. Sperka for programming the stimulus and for his help managing the experimental data. We also thank Heidi Englehardt, Xochi Navarro, Alena Druhavets and Angie Michaiel for lab support and animal training. This work was supported by the National Institute of Health (EY 10562), the Marsden Fund, and the UC Davis Research to Prevent Blindness Grant. We also thank the Training Program in Basic Neuroscience (5T32MH082174) at U.C. Davis for its support of S.W. Egger, and the Neuroscience and Cognition master’s degree program at Utrecht University for its support of S. W. Keemink.

